# A physiologically based pharmacokinetic model for CYP2E1 phenotyping via chlorzoxazone

**DOI:** 10.1101/2023.04.12.536571

**Authors:** Jonas Küttner, Jan Grzegorzewski, Hans-Michael Tautenhahn, Matthias König

## Abstract

The cytochrome P450 (CYP) superfamily of enzymes plays a critical role in the metabolism of drugs, toxins, and endogenous and exogenous compounds. The activity of CYP enzymes can be influenced by a variety of factors, including genetics, diet, age, environmental factors, and disease. Among the major isoforms, CYP2E1 is of particular interest due to its involvement in the metabolism of various low molecular weight chemicals, including alcohols, pharmaceuticals, industrial solvents, and halogenated anesthetics. Metabolic phenotyping of CYPs based on the elimination of test compounds is a useful method for assessing in vivo activity, with chlorzoxazone being the primary probe drug for phenotyping of CYP2E1. The aim of this work was to investigate the effect of changes in CYP2E1 level and activity, ethanol consumption, ethanol abstinence, and liver impairment on the results of metabolic phenotyping with chlorzoxazone. To accomplish this, an extensive pharmacokinetic dataset for chlorzoxazone was established and a physiologically based pharmacokinetic (PBPK) model of chlorzoxazone and its metabolites, 6-hydroxychlorzoxazone and chlorzoxazone-O-glucuronide, was developed and validated. The model incorporates the effect of ethanol consumption on CYP2E1 levels and activity by extending the model with a core ethanol pharmacokinetic model and a CYP2E1 turnover model. The model accurately predicts pharmacokinetic data from several clinical studies and is able to estimate the effect of changes in CYP2E1 levels and activity on chlorzoxazone pharmacokinetics. Regular ethanol consumption induces CYP2E1 over two to three weeks, resulting in increased conversion of chlorzoxazone to 6-hydroxychlorzoxazone and a higher 6-hydroxychlorzoxazone/chlorzoxazone metabolic ratio. After ethanol withdrawal, CYP2E1 levels return to baseline within one week. Importantly, liver impairment has an opposite effect, resulting in reduced liver function via CYP2E1. In alcoholics with liver impairment who also consume ethanol, these factors will have opposite confounding effects on metabolic phenotyping with chlorzoxazone.

## 1 INTRODUCTION

Cytochrome P450 (CYP) enzymes are a superfamily of heme-containing enzymes that are critical for the oxidation of drugs, toxins, and both endogenous and exogenous compounds. CYP2E1 is a major isoform that contributes approximately 20-25% of the hepatic P450 protein pool and plays an important role in the metabolism of various low molecular weight chemicals, including alcohols, drugs, industrial solvents, and halogenated anesthetics (Couto et al., 2019; Raucy et al., 1993; Tanaka et al., 2000; Zhang et al., 2016).

To assess the in vivo activity of CYP enzymes, test substances that are specifically metabolized by these enzymes can be used as probe drugs. The pharmacokinetics of the test substance and its metabolites are used to determine the function of the enzyme. Chlorzoxazone has been established as the primary probe drug for metabolic phenotyping of CYP2E1 (Bachmann and Sarver, 1996; Dreisbach et al., 1995).

Chlorzoxazone is a muscle relaxant used to treat muscle spasms and low back pain (Liv, 2012). Its primary metabolism occurs in the liver via CYP2E1-mediated 6-hydroxylation. The resulting 6-hydroxychlorzoxazone is rapidly conjugated with glucuronic acid and excreted by the kidneys, with less than 2% of the administered dose recovered in the urine as free chlorzoxazone. No unconjugated 6-hydroxychlorzoxazone is detected in blood samples, indicating that chlorzoxazone is completely metabolized and the resulting 6-hydroxychlorzoxazone is conjugated within a single pass through the liver (Liv, 2012; de Vries et al., 1994; Desiraju et al., 1983).

Metabolic phenotyping of CYP2E1 using chlorzoxazone typically involves recording plasma concentrations of chlorzoxazone and its metabolites 6-hydroxychlorzoxazone and chlorzoxazone-O-glucuronide over a period of 8 hours. To measure both 6-hydroxychlorzoxazone and its glucuronide, a glucuronidase is often used to cleave the glucuronide group and measure the concentration of 6-hydroxychlorzoxazone and its glucuronide. Various pharmacokinetic parameters can be calculated from the metabolite time courses to assess the metabolic phenotype. Alternatively, the percentage of the chlorzoxazone dose recovered in urine as CZXOGlu has been used, which ranges from 39 to 74% and shows considerable interindividual variability (Desiraju et al., 1983; de Vries et al., 1994; Dreisbach et al., 1995; Frye et al., 1998; Girre et al., 1994; Lucas et al., 1993). The reason for this incomplete recovery is not clear and could be due to incomplete absorption or the presence of alternative metabolites. Simplified phenotyping methods based on the 6-hydroxychlorzoxazone/chlorzoxazone metabolic ratio after two or four hours from a single plasma sample have been established as a proxy for CYP2E1 metabolic activity.

Studies have shown that CYP1A1 and CYP1A2, in addition to CYP2E1, may contribute to the 6-hydroxylation of chlorzoxazone in human liver microsomes (Carriere et al., 1993; Ono et al., 1995; Yamamura et al., 2015), raising concerns about whether chlorzoxazone is a suitable phenotypic probe for measuring CYP2E1 activity. Cigarette smoking is known to induce CYP1A1 and CYP1A2 (Grzegorzewski et al., 2021a); however, two studies found no effect of tobacco smoking on chlorzoxazone metabolism in vivo (Girre et al., 1994; Lucas et al., 1999). In contrast, another study using a within-subject design reported an acceleration of chlorzoxazone metabolism (Benowitz et al., 2003). These results suggest that the contribution of CYP1A1 and CYP1A2 to chlorzoxazone metabolism is small and that 6-hydroxylation of chlorzoxazone is a good marker of CYP2E1 activity.

A thorough understanding of CYP2E1 regulation is important because it plays a critical role in activating protoxins and can generate reactive oxygen species that can cause liver damage. Several factors, including genetics, diet, fasting, age, sex, environmental factors, and disease, can influence CYP2E1 activity. In particular, ethanol consumption has been shown to be a strong inducer of CYP2E1 metabolism. Several studies have shown that the liver microsomes of alcoholics have higher CYP2E1 protein levels compared to non-drinkers or non-active drinkers (Mishin et al., 1998). In addition, numerous pharmacokinetic studies have found that CYP2E1-mediated metabolism of chlorzoxazone is increased in alcoholics (Girre et al., 1994; de la Maza et al., 2000; Mishin et al., 1998; Lucas et al., 1993).

The widespread consumption of ethanol worldwide makes it a particularly interesting inducer of CYP2E1. While alcohol dehydrogenase is primarily responsible for ethanol metabolism in the liver, CYP2E1 can also metabolize ethanol, albeit with a lower affinity (about 10 mM) than alcohol dehydrogenase (0.05-1 mM) (Jiang et al., 2020). Consequently, CYP2E1 plays a role in alcohol elimination at higher ethanol concentrations, and its inducibility by ethanol contributes to metabolic tolerance in regular drinkers (Osna et al., 2017). The regulation of CYP2E1 induction by ethanol is thought to involve transcriptional and post-translational mechanisms. In human liver biopsy samples from recent drinkers, CYP2E1 mRNA levels were found to be increased compared to non-drinkers, although the transcriptional regulation of CYP2E1 is not yet fully understood (Takahashi et al., 1993). In rats, several studies have shown that ethanol can slow the degradation of CYP2E1 (Bardag-Gorce et al., 2002; Eliasson et al., 1988; Song et al., 1989). Substrate binding can block the ubiquitination site of CYP2E1, making the enzyme inaccessible to the ubiquitin-proteasome system (Banerjee et al., 2000; Roberts et al., 1995).

Chlorzoxazone-based metabolic phenotyping is widely used to evaluate alcoholic patients. However, a critical question that remains unanswered is how social or abusive alcohol consumption and abstinence from alcohol affect metabolic phenotyping via CYP2E1. Abusive alcohol use is associated with reduced liver function and a spectrum of liver disease ranging from steatosis and nonalcoholic fatty liver disease to alcoholic cirrhosis. The effect of liver disease and enzyme induction on metabolic phenotyping with chlorzoxazone remains unclear.

The aim of this study was to use a physiologically based pharmacokinetic (PBPK) approach to investigate the metabolic phenotyping of CYP2E1 using chlorzoxazone. Specifically, we aimed to address key questions regarding how changes in CYP2E1 level and activity, ethanol consumption and abstinence, and liver impairment might affect the results of metabolic phenotyping with chlorzoxazone.

## 2 MATERIAL AND METHODS

### Data

A wide range of heterogeneous data was curated for model building (parameterization) and subsequent model validation (comparison of model predictions with clinical data). We systematically searched PubMed using the search string “(chlorzoxazone) AND (pharmacokinetics) AND ((CYP2E1) OR (P4502E1))”. The result set of publications was searched for pharmacokinetic time course data and pharmacokinetic parameters of chlorzoxazone. Studies reporting pharmacokinetic data in healthy and/or alcoholic subjects were of particular interest. The initial corpus of publications was expanded based on references in the initial set of publications. From the selected studies, information on the subjects and groups (e.g. age, sex, disease, medication), the route of administration and dose of chlorzoxazone, and the pharmacokinetics of chlorzoxazone were manually curated. Established data curation workflows for pharmacokinetic data were applied to digitize data from figures, tables, and text (Grzegorzewski et al., 2021a). In addition data on the pharmacokinetics of ethanol were curated from a single study identified by searching for “(ethanol) AND (pharmacokinetics)” and manual screening of the results (Wilkinson et al., 1977). All data are available in the pharmacokinetics database PK-DB (https://pk-db.com) (Grzegorzewski et al., 2021b). An overview of the 29 studies with their respective chlorzoxazone and ethanol dosing protocols is provided in Tab. 1.

**Table 1:**
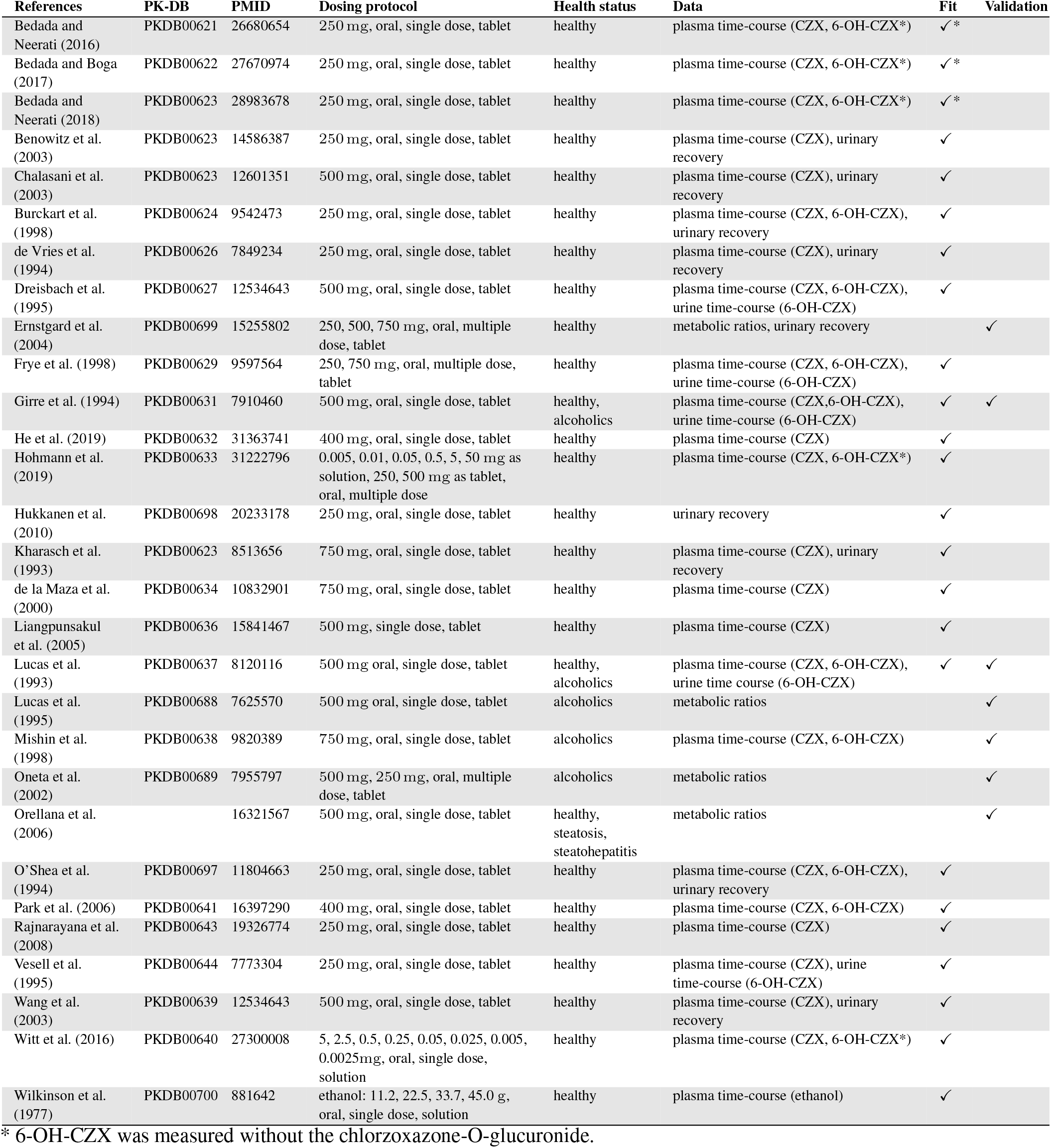
Overview of curated clinical studies.

### Model

The model was encoded in the Systems Biology Markup Language (SBML) (Hucka et al., 2019; Keating et al., 2020) and developed using sbmlutils (Kö nig, 2021b), a collection of Python utilities for building SBML models, and cy3sbml (Kö nig et al., 2012; Kö nig and Rodriguez, 2019), a visualization software for SBML. The model is an ordinary differential equation (ODE) model which is numerically solved by sbmlsim (König, 2021a) based on the high-performance SBML simulator libroad-runner (Somogyi et al., 2015; Welsh et al., 2023). The model is available under the CC-BY 4.0 license at https://github.com/matthiaskoenig/chlorzoxazone-model. Version 0.9.2 was used in this paper (Küttner and König, 2023).

#### PBPK model of chlorzoxazone and ethanol

The physiologically based pharmacokinetic (PBPK) model is hierarchically organized (Fig. 1) and allows simulation of the time courses of chlorzoxazone, 6-hydroxychlorzoxazone, chlorzoxazone-O-glucuronide, and ethanol. The top layer represents the whole body and systemic circulation coupled with organ models for lung, liver, kidney, intestine, and the rest compartment. Organs not relevant to chlorzoxazone metabolism are included in the rest compartment. The metabolic and transport reactions for chlorzoxazone and its metabolites are included in the organ models.

**Figure 1.**
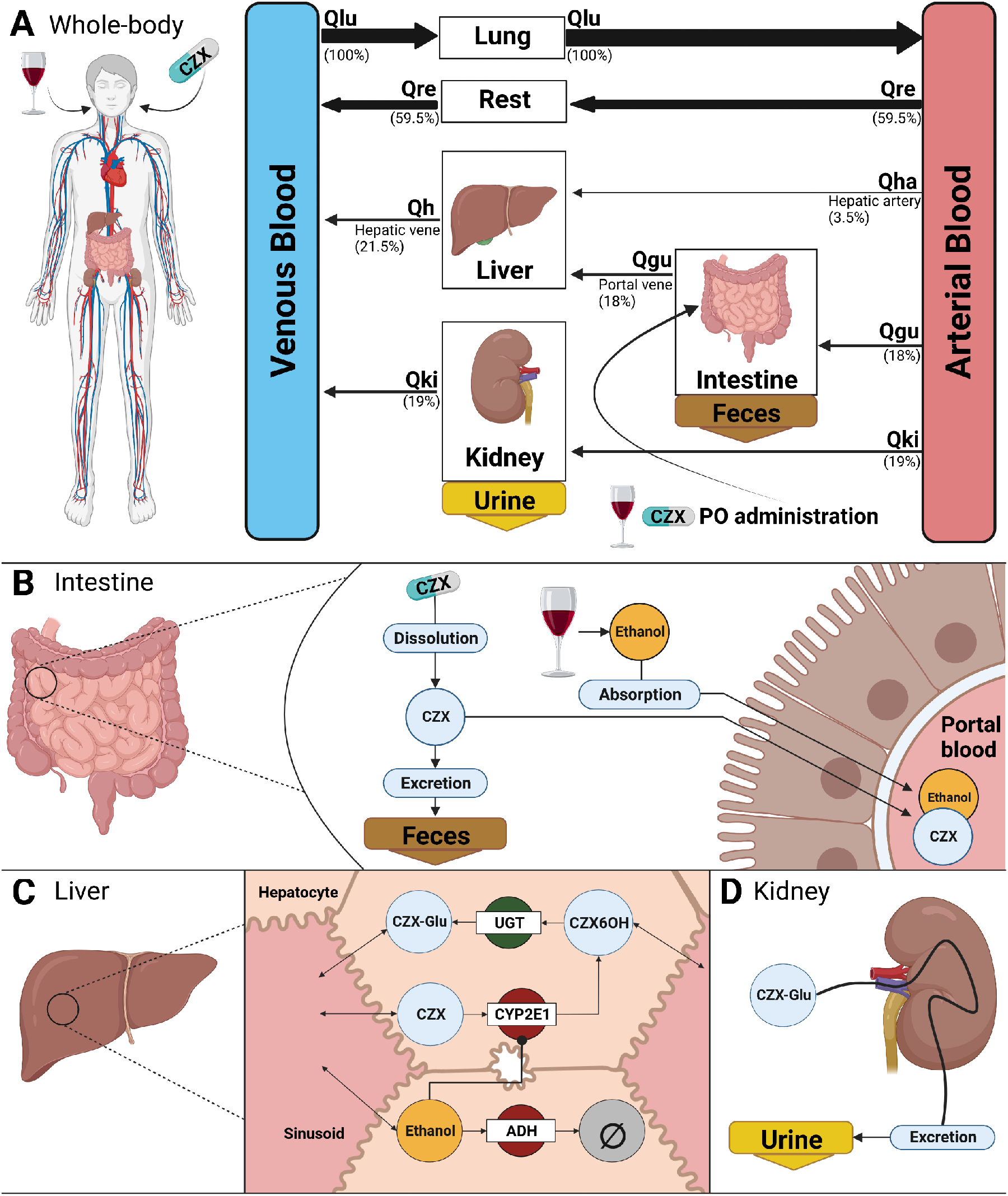
Human PKPK model of chlorzoxazone and ethanol. **(A)** Whole body model consisting of liver, kidney, lung and rest compartment. Organs with minor relevance for chlorzoxazone are not modelled explicitly and are condensed in the rest compartment. The organs are connected *via* the systemic circulation, denoted by arrows. The arrow widths are proportional to the relative blood flow through the corresponding route. chlorzoxazone and ethanol are administered orally. **(B)** Intestine model consisting of dissolution, absorption and excretion for chlorzoxazone. The dissolution depends on the application form (tablet, oral solution) and determines how fast chlorzoxazone becomes available for absorption. Only a fraction of the dose is absorbed into the systemic circulation, with the remainder being excreted into the feces. Administered ethanol is instantly available for absorption and fully absorbed. **(C)** Liver model for the conversion chlorzoxazone and the elimination of ethanol. chlorzoxazone is converted to 6-hydroxychlorzoxazone mediated by CYP2E1. 6-hydroxychlorzoxazone is glucuronidated by UGT. Ethanol is eliminated by the alcohol dehydrogenase. Ethanol metabolism by CYP2E1 is neglected. CYP2E1 is induced by the presences of ethanol. **(D)** Kidney model of urinary excretion of chlorzoxazone-O-glucuronide. Created with BioRender.com

Transport reactions describe the import and export of chlorzoxazone, its metabolites, and ethanol between plasma and organ compartments. Transport reactions are modeled by mass action kinetics of the form *v* = *k*_i_ *·* (*c*_e_ *− c*_i_ *· f*), where *k*_i_ is the import rate constant, *f* is a factor that scales the import rate to achieve an equilibrium tissue distribution, and *c*_e_ and *c*_i_ are the plasma and compartment concentrations, respectively.

All metabolic reactions of chlorzoxazone ta*k*e place in the liver compartment. Chlorzoxazone is converted to 6-hydroxychlorzoxazone (6-hydroxylation), 6-hydroxychlorzoxazone is converted to chlorzoxazone-O-glucuronide (glucuronidation), and ethanol is eliminated. All metabolic reactions are modeled using irreversible Michaelis-Menten kinetics of the form 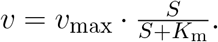

#### Ethanol induction of CYP2E1

A protein stabilization model was implemented to describe the induction of CYP2E1 by ethanol. CYP2E1 was implemented as a dimensionless quantity (relative amount) that is produced and degraded at rates *k*_p_ and *k*_d_, respectively. The degradation rate of CYP2E1 was set based on the reported half-life of CYP2E1 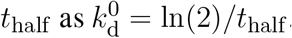. A steady state concentration of 1 was obtained by assuming *k*_p_ = *k*_d_.

The amount of CYP2E1 modulates the v_max_ value of the 6-hydroxylation reaction catalyzed by CYP2E1. Ethanol inhibits the degradation of CYP2E1, resulting in an increase in CYP2E1. The inhibition was modeled by 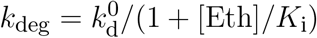 where 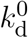 is the degradation rate in the absence of alcohol, [Eth] is the ethanol concentration, and K_i_ is the inhibition constant of ethanol on the degradation.

### Model parametrization

Values for organ volumes and tissue blood flows were taken from the literature (ICRP, 2002; Jones and Rowland-Yeo, 2013). Eight model parameters were fitted for the chlorzoxazone metabolism model and four parameters were fitted for the ethanol metabolism model. The parameters were determined by minimizing the residuals between model predictions and time-course data.

A large subset of the curated clinical data was used to parameterize the chlorzoxazone model. This data included time course data for plasma concentrations of chlorzoxazone, 6-hydroxychlorzoxazone, the sum of 6-hydroxychlorzoxazone and chlorzoxazone-O-glucuronide, and urinary levels of chlorzoxazone-O-glucuronide. It covered a wide range of chlorzoxazone doses from 0.005 mg to 750 mg and included data for chlorzoxazone administration by tablet and oral solution. The data used to parameterize the model were selected according to the following criteria Subjects were healthy and chlorzoxazone was administered exclusively, i.e., no co-administration of other drugs or cocktail administrations. The ethanol elimination model was parameterized using a single study that provided ethanol time course data for four doses (Wilkinson et al., 1977). The optimization problem was solved using SciPy’s least-squares method and differential evolution algorithm (Virtanen et al., 2020). For the objective cost function F, which depends on the parameters 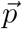, a simple L2 norm consisting of the sum of weighted residuals was used.

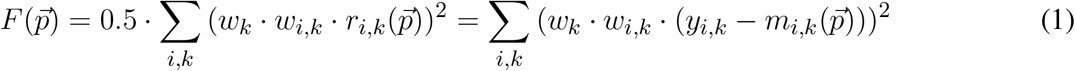

where 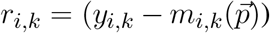 is the residual of time point i in time course *k* for model prediction 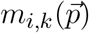 and the corresponding data point *y*_*i,k*_; *w*_*i,k*_ is the weighting of the respective data point i in time course *k* based on the error of the data point and *w*_*k*_ = the weighting factor of time course *k*. Weighting of time courses was based on the number of subjects per study. The data used for the parameter fit is listed in Tab. 1. The final parameter set given in Tab. 2 was determined using 10 runs of the local least squares optimization.

**Table 2:** Overview of model parameters.

| Physiological parameter | Description | Value | Unit | Reference |
| --- | --- | --- | --- | --- |
| BW | body weight | 75 | kg | ICRP (2002) |
| HEIGHT | body height | 170 | cm | ICRP (2002) |
| COBW | cardiac output per body weight | 1.548 | ml/s/kg | ICRP (2002); de Simone et al. (1997) |
| HCT | hematocrit | 0.51 |  | Vander (2001); Herman (2016) |
| f_lumen | fraction lumen of intestine | 0.9 |  |  |
| FVgu | gut fractional tissue volume | 0.0171 | l/kg | Jones and Rowland-Yeo (2013); ICRP (2002) |
| FVli | liver fractional tissue volume | 0.0021 | l/kg | Jones and Rowland-Yeo (2013); ICRP (2002) |
| FVlu | lung fractional tissue volume | 0.076 | l/kg | Jones and Rowland-Yeo (2013); ICRP (2002) |
| FVve | venous fractional tissue volume | 0.0514 | l/kg | Jones and Rowland-Yeo (2013); ICRP (2002) |
| FVar | arterial fractional tissue volume | 0.0257 | l/kg | Jones and Rowland-Yeo (2013); ICRP (2002) |
| FVpo | portal fractional tissue volume | 0.001 | l/kg | Jones and Rowland-Yeo (2013); ICRP (2002) |
| FVhv | hepatic venous fractional tissue volume | 0.001 | l/kg |  |
| FVgu | gut fractional tissue blood flow | 0.018 | l/kg | Jones and Rowland-Yeo (2013) |
| FQki | kidney fractional tissue blood flow | 0.19 |  | Jones and Rowland-Yeo (2013) |
| FQh | hepatic fractional tissue blood flow | 0.215 |  | Jones and Rowland-Yeo (2013) |
| Mr_czx | molecular weight CZX | 169.56 | g/mole | pubchem.compound/2733 |
| Mr_czx6oh | molecular weight 6-OH-CZX | 185.56 | g/mole | pubchem.compound/2734 |
| Mr_czxoglu | molecular weight CZX-O-Glu | 361.69 | g/mole | pubchem.compound/129522086 |
| Mr_eth | molecular weight ethanol | 46.07 | g/mole | pubchem.compound/702 |
| LI_Ki_cyp2e1_deg_ethanol | Ethanol-CYP2E1 inhibition constant | 0.01 | μM | adjustment by eye |
| LI_cyp2e1_thalf | half-time of CYP2E1 | 1.8 | day | adjustment by eye |
| Fit parameter | Description | Value | Unit |  |
| Ka_dis_tablet_czx | dissolution rate of chlorzoxazone tablet | 0.521 | 1/hr |  |
| GU_Ka_abs_czx | rate of gut chlorzoxazone absorption | 1.325 | 1/hr |  |
| GU_F_abs_czx | bio-availability for chlorzoxazone absorption | 0.593 | dimensionless |  |
| LI_CZXOX_Vmax | v<sub>max</sub> for chlorzoxazone 6-hydroxylation | 7.527 | μmol/min/l |  |
| LI_CZXOX_Km | K<sub>M</sub> value for 6-hydroxylation of chlorzoxazone | 18.929 | μM |  |
| LI_CZXOGLU_Vmax | v<sub>max</sub> for glucuronidation of 6-OH-CZX | 11.346 | μmole/min/l |  |
| LI_CZXOGLU_Km | K<sub>M</sub> for glucuronidation of 6-OH-CZX | 0.255 | μM |  |
| KI_CZXOGLU_EX_k | urinary excretion rate of CZX-O-Glu | 2.611 | 1/min |  |
| GU_Ka_abs_eth | absorption rate of ethanol | 2.59 | 1/hr |  |
| LI_ETHEL_Vmax | v<sub>max</sub> for ethanol elimination | 0.033 | mmole/min/kg |  |
| LI_ETHEL_KM | K<sub>M</sub> for ethanol elimination | 0.046 | mM |  |
| Kp_ethanol | ethanol tissue partition coefficient | 0.59 | dimensionless |  |
| Scan parameter | Description | Value | Unit | Range |
| LI_f_cyp2e1 | scaling factor of CYP2E1 vactivity | 1 | dimensionless | 0.1 - 100 |
| LI_CZXOX_Km | CYP2E1 K<sub>M</sub> value | 18.929 | μM | 1.8929 - 189.29 |

### Pharmacokinetics parameters

Pharmacokinetic parameters of chlorzoxazone were calculated from the plasma-concentration time courses and urinary excretion using standard non-compartmental methods (Urso et al., 2002). The elimination rate k_el_ [1/min] was calculated via linear regression in logarithmic space in the decay phase. The area under the curve AUC [mmole*·*min/L] was calculated via the trapezoidal rule and interpolated to infinity. Clearance Cl [ml/min] was calculated as *Cl = k*_*el*_ *· V*_*d*_ with the apparent volume of distribution V_*d*_ *as V d = D/(AUC*_*∞*_ *· k*_*el*_). D is the applied dose of chlorzoxazone.

## 3 RESULTS

### 3.1 Database of chlorzoxazone pharmacokinetics

To parameterize and validate the chlorzoxazone pharmacokinetic model, we curated a data set consisting of 29 clinical trials. Most of the studies investigated drug-drug interactions, the effect of lifestyle and physiological conditions such as alcoholism, obesity or diabetes, and the effect of different doses. In all studies, chlorzoxazone was administered orally, mostly as a tablet and in rare cases as an oral solution. The standard doses given were 250, 500 or 750 mg. In some cases, 400 mg was administered. In two dose escalation studies, the doses ranged from 0.05 mg to 500 mg. In most studies, plasma concentrations were measured for chlorzoxazone (n=23) and the metabolite 6-hydroxychlorzoxazone (n=13). The plasma concentration of 6-hydroxychlorzoxazone was reported either as the concentration of unconjugated 6-hydroxychlorzoxazone (n=4) or as the sum of unconjugated 6-hydroxychlorzoxazone and the glucuronide chlorzoxazone-O-glucuronide (n=9). Some studies (n=5) also reported the time course of chlorzoxazone-O-glucuronide recovered in urine, either as the amount or as the percentage of the administered dose recovered.

### 3.2 PBPK model of chlorzoxazone

We developed a physiologically-based pharmacokinetics (PBPK) model for chlorzoxazone coupled with a model of CYP2E1 regulation by ethanol (Fig. 1) using the curated data. The model is hierarchically organized, with the top layer representing the whole body comprising the lung, liver, kidney, intestine, and rest compartment, and transport via the systemic circulation. The intestine model (Fig. 1B) describes the dissolution, absorption, and excretion of chlorzoxazone. Only a fraction of the administered chlorzoxazone dose is absorbed into the systemic circulation, with the remainder being excreted in the feces. In the liver (Fig. 1C), chlorzoxazone is metabolized to 6-hydroxychlorzoxazone via CYP2E1 and subsequently converted to chlorzoxazone-O-glucuronide. The kidney model describes the renal excretion of chlorzoxazone-O-glucuronide into the urine. Ethanol is absorbed into the systemic circulation (Fig. 1B) in the intestine and eliminated in the liver (Fig. 1C). Ethanol affects chlorzoxazone metabolism by inhibiting the degradation of CYP2E1 in the liver, resulting in increased CYP2E1 and conversion of chlorzoxazone. The mathematical details of the model are described in the Materials and Methods.

### 3.3 Model performance

The model accurately predicts pharmacokinetic data from various clinical studies (Fig. 2). Our model successfully described the concentration profiles of chlorzoxazone, 6-hydroxychlorzoxazone, chlorzoxazone-O-glucuronide, and the amount of chlorzoxazone-O-glucuronide excreted in the urine over time. The model was able to predict these profiles for doses ranging from 0.005 mg to 750 mg administered as tablets or oral solutions in the studies we used for model parameterization (Bedada and Neerati, 2016; Bedada and Boga, 2017; Bedada and Neerati, 2018; Burckart et al., 1998; de Vries et al., 1994; Dreisbach et al., 1995; Frye et al., 1998; Girre et al., 1994; He et al., 2019; de la Maza et al., 2000; Liangpunsakul et al., 2005; Lucas et al., 1993; Park et al., 2006; Rajnarayana et al., 2008; Vesell et al., 1995; Wang et al., 2003; Hohmann et al., 2019; Witt et al., 2016). Refer to Tab. 1 for details on the respective doses and application forms.

**Figure 2.**
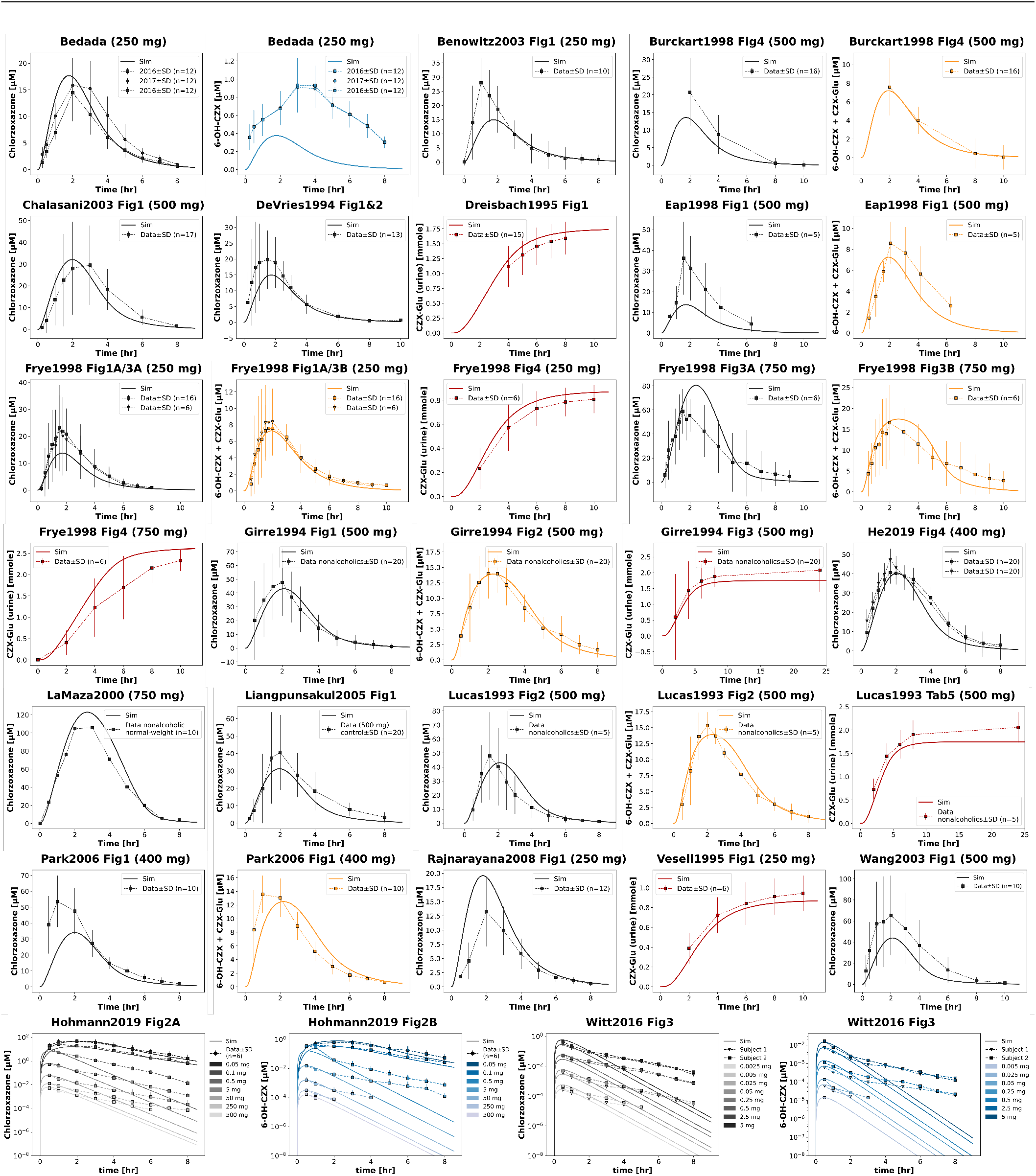
Performance of the chlorzoxazone model. The model simulates time courses for the plasma concentrations of chlorzoxazone, chlorzoxazone-O-glucuronide, 6-hydroxychlorzoxazone and the urinary amounts of chlorzoxazone-O-glucuronide. The time-courses shown in this plot were used for parameter fitting, except for the 6-hydroxychlorzoxazone time-courses from the Bedada studies. Studies were selected when they met the following criteria: (1) only chlorzoxazone was administered (no cocktail or co-administrations) (2) the subjects are adults (3) data for more than one subject was reported. Data from (Bedada and Neerati, 2016; Bedada and Boga, 2017; Bedada and Neerati, 2018; Benowitz et al., 2003; Burckart et al., 1998; Chalasani et al., 2003; Dreisbach et al., 1995; Eap et al., 1998; Girre et al., 1994; He et al., 2019; de la Maza et al., 2000; Liangpunsakul et al., 2005; Lucas et al., 1993; Park et al., 2006; Rajnarayana et al., 2008; Vesell et al., 1995; Wang et al., 2003; Hohmann et al., 2019; Witt et al., 2016).

To investigate the dose-dependency of pharmacokinetic parameters, we analyzed the c_max_ of chlorzoxazone and 6-hydroxychlorzoxazone, as well as the metabolic ratio at two hours for different doses per body weight. Our model predicted an increase in both chlorzoxazone and 6-hydroxychlorzoxazone c_max_ with increasing dose per body weight, and a decrease in the metabolic ratio at two hours. The predicted dose-dependency was generally consistent with the observed pharmacokinetic data (Fig. 3).

**Figure 3.**
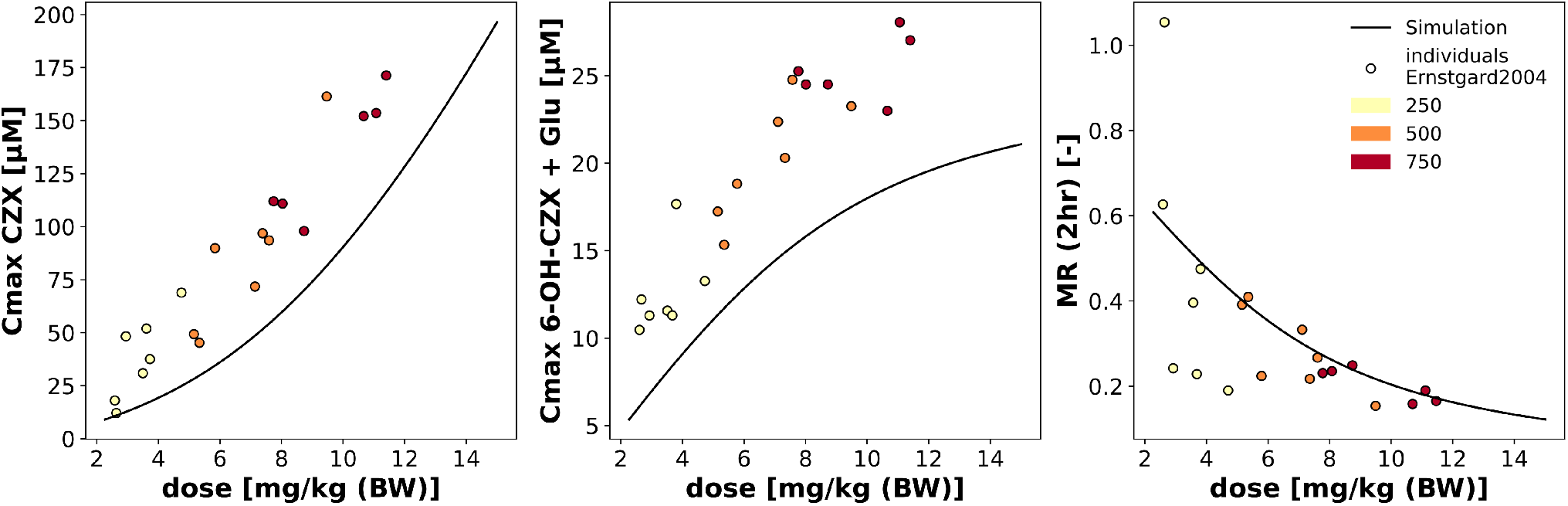
Prediction of pharmacokinetics based on body weight normalized dose. The model body weight was scanned linearly from 50 to 110 kg for a doses of 250, 500, and 750 mg. For each time-course, cmax of chlorzoxazone, 6-hydroxychlorzoxazone + chlorzoxazone-O-glucuronide, and the metabolic ratio at 2hr, were calculated. The results were sorted by the dose normalized by body weight (black solid lines). The simulation was compared to individual data from (Ernstgard et al., 2004)

### Effect of CYP2E1 on chlorzoxazone pharmacokinetics

To explore the impact of changes in CYP2E1 activity and affinity on the conversion of chlorzoxazone to 6-hydroxychlorzoxazone in metabolic phenotyping, we examined the effects of variations in *v*_*max*_ and K_M_ by systematically scanning both parameters (Fig. 4). The time courses of chlorzoxazone and its metabolites at a dose of 250 mg for *v*_*max*_ and *K*_*M*_ are shown in the left columns of Fig. 4A and B, respectively. As CYP2E1 activity (*v*_*max*_) increases above the reference value, chlorzoxazone plasma concentrations decrease, while 6-hydroxychlorzoxazone, 6-hydroxychlorzoxazone + chlorzoxazone-O-glucuronide and urinary recovery show minimal change. Conversely, decreasing CYP2E1 leads to increased plasma concentrations of chlorzoxazone and reduced 6-hydroxychlorzoxazone, 6-hydroxychlorzoxazone + chlorzoxazone-O-glucuronide during the first hours, followed by an increase after about 5 hours, with a decrease in urinary recovery. The variation in CYP2E1 affinity (*K*_*M*_) shows opposite effects.

**Figure 4.**
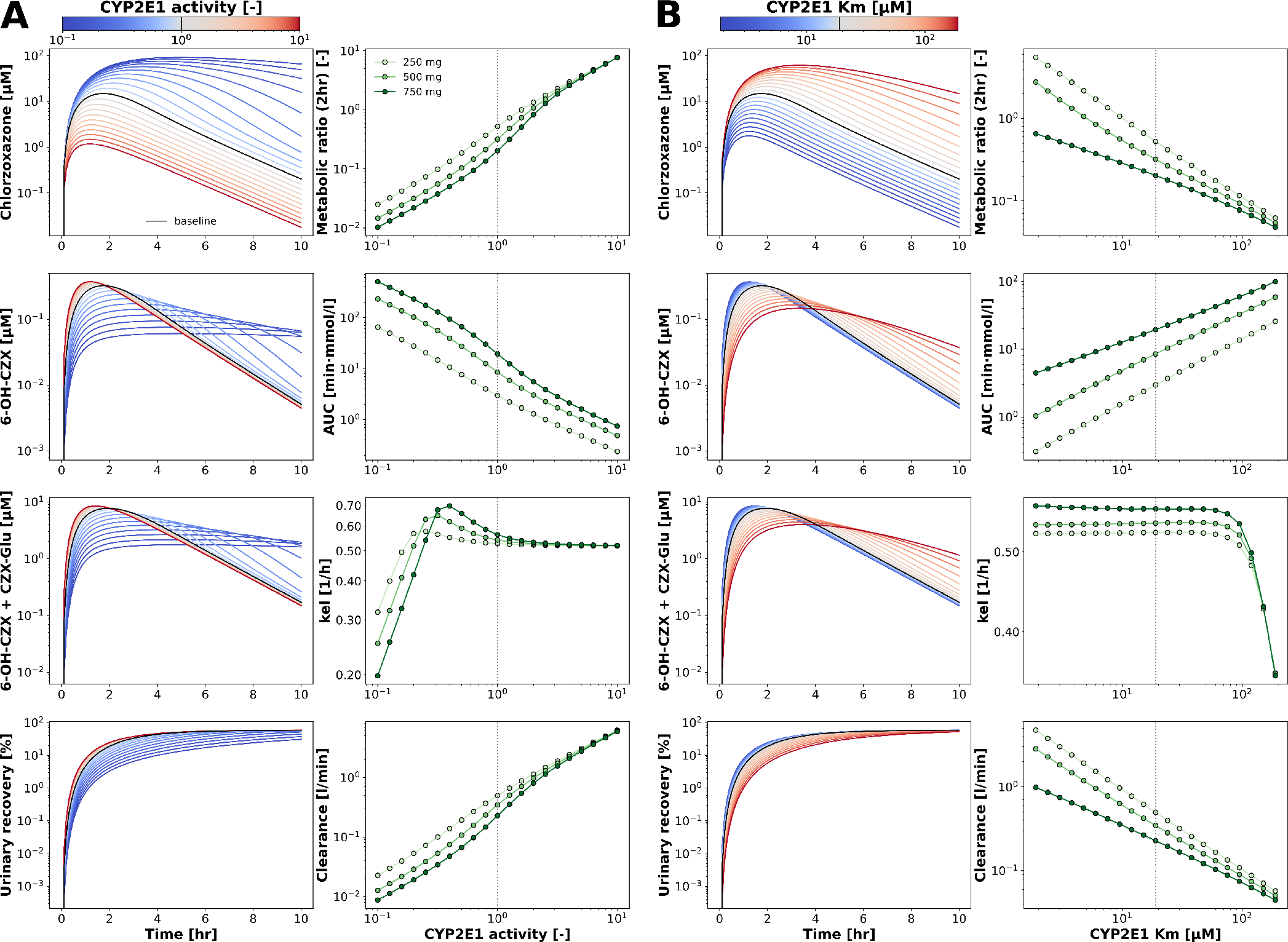
Parameter scans (A, B) Scan for the CYP2E1 activity and the K_M_ value, respectively. The parameters were scanned using a log range from (−1, 1) which was multiplied with the fitted reference value of the scanned parameter (CYP2E1 activity, 1; CYP2E1 K_M_, 18.929 µM). The scan was repeated for three doses (250, 500, 750 mg), and the pharmacokinetic parameters metabolic ratios (2 hr), area under the curve (AUC), elimination rate (kel), and clearance were calculated. The time-courses for the 250 mg dose are shown on the left column, and the pharmacokinetic parameters in the right column.

The scan was performed for 250, 500, and 750 mg chlorzoxazone doses to evaluate the influence of different chlorzoxazone doses. pharmacokinetic parameters were calculated for each time course. The 2-hour metabolic rate (MR) and chlorzoxazone clearance increase with increasing CYP2E1 v_max_, while the AUC decreases. In contrast, the K_M_ of CYP2E1 shows an inverse relationship for these pharmacokinetic parameters. The elimination rate kel is not significantly affected by either v_max_ or K_M_.

As expected, both changes in CYP2E1 activity (*v*_*max*_) and affinity (*K*_*M*_) have a significant impact on chlorzoxazone pharmacokinetics and metabolic phenotyping results based on chlorzoxazone.

### 3.5 CYP2E1 induction in alcoholic subjects

Subsequently, we aimed to determine whether the observed changes in chlorzoxazone pharmacokinetics and metabolic phenotyping results in alcoholics could be attributed to increased CYP2E1 activity resulting from induction of CYP2E1 protein levels (Mishin et al., 1998). To determine whether our model could qualitatively replicate the pharmacokinetic changes observed in alcoholic subjects, we compared a parameter scan for CYP2E1 activity with time course data from two clinical studies (Girre et al., 1994; Lucas et al., 1993). Both studies contrasted a group of alcoholic subjects with a group of non-drinking controls. A 500 mg dose of chlorzoxazone was administered, and plasma concentrations of chlorzoxazone, 6-hydroxychlorzoxazone, and chlorzoxazone-O-glucuronide, as well as urinary recovery, were measured for at least 8 hours. The differences observed in both studies are consistent with the predictions of the pharmacokinetic model (Fig. 5). In the alcoholic groups, the maximum concentrations achieved for chlorzoxazone are lower because elevated CYP2E1 levels accelerate chlorzoxazone hydroxylation. Despite the significant differences in the chlorzoxazone curves, the maxima of the 6-hydroxychlorzoxazone curves show only a modest increase. The urine recovery curves show similarities between the alcoholic and control groups.

**Figure 5.**
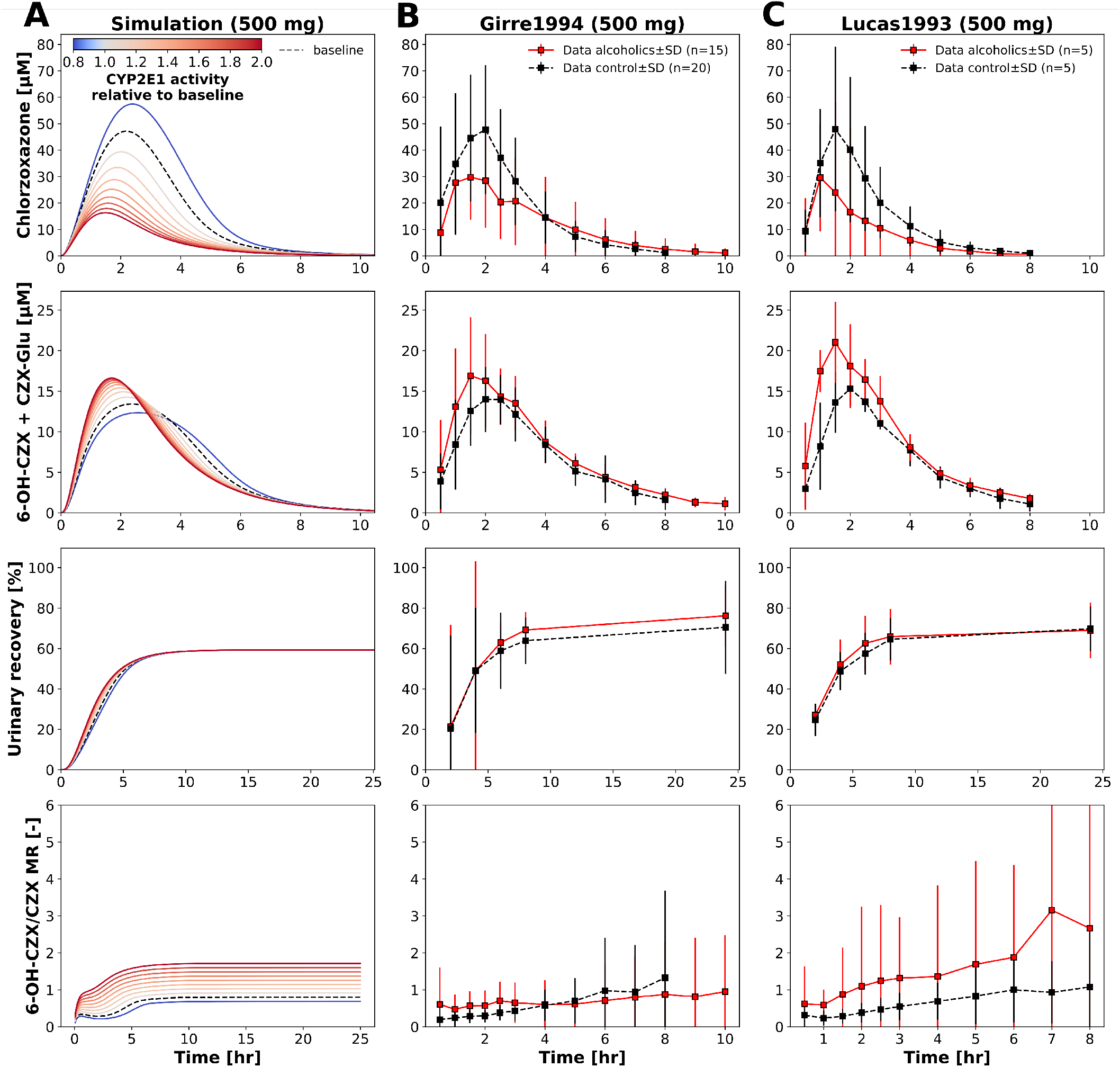
CYP2E1 induction in alcoholic subjects. **(A)** Parameter scan of the CYP2E1 activity. Simulated time-courses are shown, for plasma chlorzoxazone, chlorzoxazone-O-glucuronide, urinary recovery and metabolic ratio (2 hr). The black dashed line denotes the time-course belonging to the fitted baseline activity. **(B, C)** Time course data from (Girre et al., 1994; Lucas et al., 1993) for non-drinking control (black dashed line) and alcoholic subjects (red solid line). The metabolic ratios were calculated from the time-course data of chlorzoxazone and chlorzoxazone-O-glucuronide. The alcoholic subjects form (Girre et al., 1994) consumed (333 *±* 191) g (mean *±* SD) of alcohol over (15.7 *±* 10.7) No data about alcohol amounts consumed were reported by (Lucas et al., 1993). Both studies administered 500 mg of chlorzoxazone.

The 2-hour metabolic ratio (MR) is higher in alcoholics compared to controls. In conclusion, changes in CYP2E1 activity resulting from the induction of CYP2E1 could account for the observed alterations in chlorzoxazone pharmacokinetics in alcoholic patients.

### 3.6 Changes in pharmacokinetic parameters due to changes in microsomal CYP2E1

While numerous studies have indicated that in vitro CYP2E1 metabolism correlates with CYP2E1 quantity in human liver microsomes, only a handful of investigations have examined the relationship between CYP2E1 amount and in vivo chlorzoxazone pharmacokinetics. To assess whether microsomal protein quantity could serve as a proxy for CYP2E1 activity, we simulated two studies (Mishin et al., 1998; Orellana et al., 2006) that reported the dependence of chlorzoxazone-O-glucuronide c_max_ and the 2-hour MR on microsomal protein concentration, respectively. We normalized the reported protein concentration to the mean of the abstinent group and the control for Mishin et al. (1998) and Orellana et al. (2006), respectively.

In line with the data, c_max_ rises with increasing CYP2E1 activity. However, our model does not adequately capture the overall increase of chlorzoxazone-O-glucuronide c_max_ reported by Mishin et al. (1998), as it predicts saturated 6-hydroxychlorzoxazone and chlorzoxazone-O-glucuronide concentrations with an induction of CYP2E1 activity (Fig. 6A). Conversely, the correlation between MR and microsomal protein concentrations reported byOrellana et al. (2006) for the control, steatosis, and steatohepatitis groups was in excellent agreement with the model (Fig. 6B).

**Figure 6.**
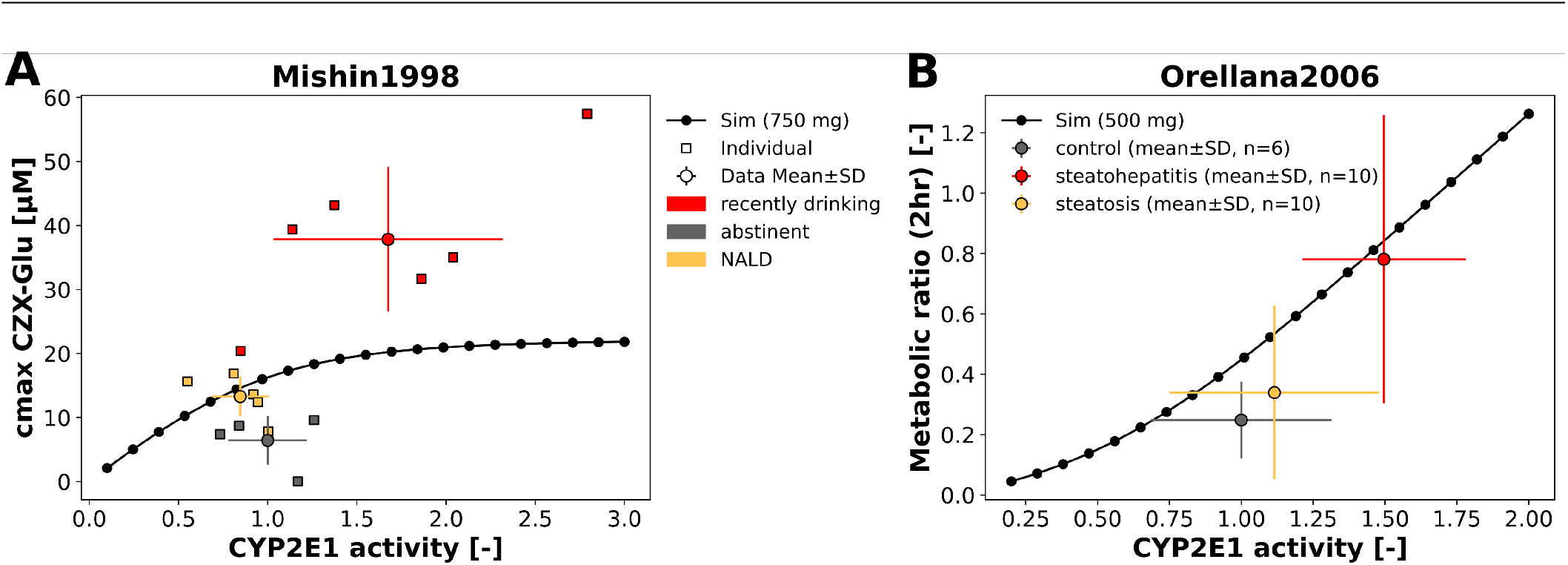
Predicted pharmacokinetics based in microsomal CYP2E1 concentrations. **(A)** Model prediction of the maximum concentration of plasma chlorzoxazone-O-glucuronide compared to data of (Mishin et al., 1998). **(B)** Model prediction of the metabolic ratio (2 hr) compared to data from (Orellana et al., 2006). The CYP2E1 activity parameter was scanned and the respective pharmacokinetic parameter was calculated. The microsomal protein concentrations reported by the studies were normalized so that the value of the abstinent (Mishin et al., 1998) or control group (Orellana et al., 2006) match the CYP2E1 activity of (`LI_f_cyp2e1` = 1).

### 3.7 Effect of chronic ethanol consumption on the induction of CYP2E1 and CZX pharmacokinetics

Our model demonstrates that the accelerated metabolism of chlorzoxazone in induced subjects can be attributed to an increase in CYP2E1 activity. Additionally, we explored the effects of chronic alcohol consumption on CYP2E1 over time. To achieve this, we simulated a three-phase dosing protocol designed to examine the impact of prolonged, moderate alcohol intake on CYP2E1 induction. In the first phase (pre-drinking), lasting five days, no ethanol was administered. During the second phase (drinking), 40 g of ethanol was administered daily at 8 pm for 30 days. In the final phase (withdrawal), no ethanol was provided (Fig. 7). Throughout the simulation experiment, 500 mg of chlorzoxazone was administered every day at 8 am. We employed this model to predict data from clinical studies that investigated comparable experimental protocols (Lucas et al., 1995; Mishin et al., 1998; Oneta et al., 2002). The previously described simulation experiment was tailored to align with the protocols used in the respective studies, taking into account factors such as dosage and physiological data of the subject groups, if available.

**Figure 7.**
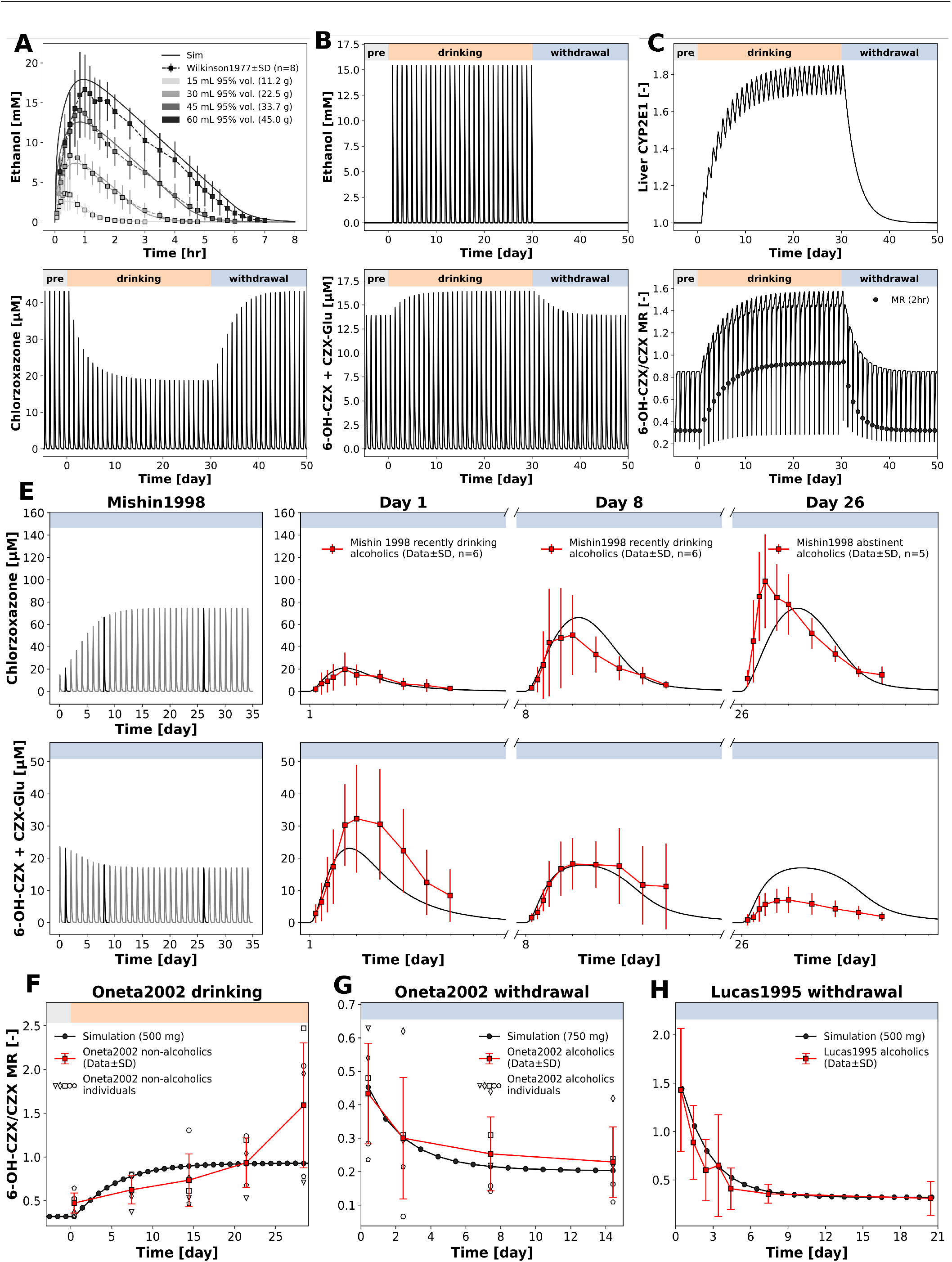
Effect of chronic ethanol consumption on the induction of CYP2E1 and chlorzoxazone pharmacokinetics. We coupled the alcohol metabolism model with the chlorzoxazone pharmacokinetic model to study co administration of alcohol and chlorzoxazone over a time of multiple weeks. **(A)** The alcohol metabolism model was parametrized using data from (Wilkinson et al., 1977). The simulation experiment consists of three phases, The pre-drinking, drinking and withdrawal phase. Over all three phases a single chlorzoxazone dose was administered at 8:00 am. During the drinking phase, a single oral dose of 40 g alcohol was administered at 8pm (**B**). The administration of alcohol causes an increase of the CYP2E1 level over time (**D**) and thereby affects the chlorzoxazone pharmacokinetics (**D**). Several clinical studies investigated the change in chlorzoxazone pharmacokinetics during the start of drinking (Oneta et al. (2002)) and withdrawal (Mishin et al. (1998); Lucas et al. (1995); Oneta et al. (2002)). We used the model to simulate those studies in silico. For the simulation of the withdrawal studies Mishin1998 (**E**), Oneta2002 (**G**), and Lucas1995 (**H**), the model was initialized with an initial CYP2E1 amount (`LI_cyp2e1`), such that the simulated time point (or time-course for chlorzoxazone for (Mishin et al., 1998)) at day 1 matched the corresponding data point. The initial CYP2E1 activity values were 3.0, 2.7 and 1.575 for Mishin1998, Lucas1995, and Oneta2002, respectively. The CYP2E1 half-life (`LI_cyp2e1_thalf`) was determined to be 1.8 d. For the simulation of the drinking group (Oneta et al. (2002)) the protocol, described above, was used. The inhibition constant (`LI_Ki_cyp2e1_deg_ethanol`) that determines how strongly alcohol inhibits the degradation of CYP2E1 was set to 0.01 µM.

The conducted experimental studies either focused on the withdrawal phase (Lucas et al., 1995; Mishin et al., 1998; Oneta et al., 2002) or the drinking phase (Oneta et al., 2002). For the withdrawal phase, recently drinking alcoholics were tested with chlorzoxazone at multiple time points after their last drink. Consequently, we initialized the corresponding simulation experiment with a CYP2E1 level that aligned with the first measurement point reported in the clinical studies. We identified varying values for the initial CYP2E1 amount, corresponding to a 3-fold, 2.7-fold, and a 1.575-fold CYP2E1 induction for Mishin1998, Lucas1995, and Oneta2002, respectively. The initial CYP2E1 amount decayed exponentially over the course of the experiment. By adjusting the CYP2E1 half-life to ensure the model output curve matched the reported data, we determined a half-life of CYP2E1 induction of approximately 2 days (Fig. 7E, G, H). Only one clinical study reported the dynamics of CYP2E1 induction in subjects who recently began drinking(Oneta et al., 2002). We simulated this study using the drinking phase of the previously described protocol and the CYP2E1 half-life determined from the withdrawal simulation experiments. The model predicts an increase in the metabolic ratio that plateaus after 2 weeks (Fig. 7F). However, the data from Oneta et al. (2002) show no saturation of induction after 4 weeks.

In summary, our model effectively predicts the changes in chlorzoxazone pharmacokinetics and metabolic phenotyping due to alcohol induction and withdrawal, showing good agreement with the data.

### 3.8 Compensatory effect of CYP2E1 induction and cirrhosis on metabolic phenotyping

Chronic heavy drinking can lead to alcoholic liver disease, which can progress to alcoholic liver cirrhosis. To assess the effects of cirrhosis in combination with CYP2E1 induction as observed in alcoholics on liver function, we performed a parameter scan, scanning the CYP2E1 activity (`LI_f_cyp2e1`) over a range from 0.5 to 4.0 and the cirrhosis parameter (`f_cirrhosis`) of our model for four cirrhosis grades: control (`f_cirrhosis` = 0), mild cirrhosis (`f_cirrhosis`=0.40, CTP A), moderate cirrhosis (`f_cirrhosis`=0.70, CTP B), and severe cirrhosis (`f_cirrhosis`=0.81, CTP C) as previously described (Köller et al., 2021a,b).

The cirrhosis parameter in our model determines the fraction by which the volume of the liver, as well as the blood flow through the liver, is reduced. Because of this two-factor reduction in liver function, the cirrhosis grade exhibits a non-linear effect on chlorzoxazone pharmacokinetics. To assess the actual CYP2E1 activity in a cirrhotic liver, we calculated the effective CYP2E1 activity, which is determined by 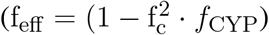, where *f*_*c*_ is the cirrhosis grade, and *f*_CYP_ denotes the CYP2E1 activity. The effective CYP2E1 activity directly translates to the predicted pharmacokinetic parameter (Fig. 8A).

**Figure 8.**
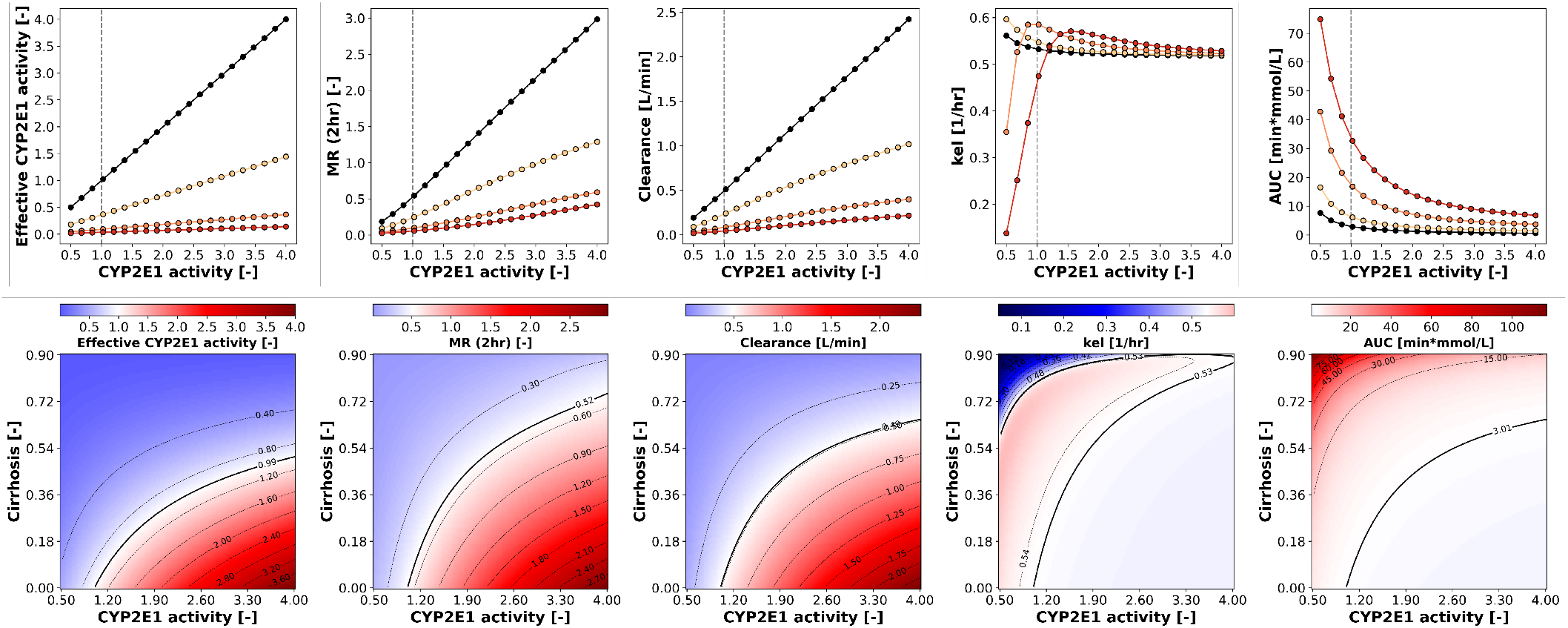
Combined effect of CYP2E1 induction and Cirrhosis. **(A)** Effect of the CYP2E1 activity on pharmacokinetic parameters for 4 different cirrhosis grades. A parameter scan for CYP2E1 (`LI_f_cyp2e1`) activity was performed for four Cirrhosis grades: Control (`f_cirrhosis` = 0), mild Cirrhosis (`f_cirrhosis` = 0.40), moderate Cirrhosis (`f_cirrhosis` = 0.70), severe Cirrhosis (`f_cirrhosis` = 0.81). **(B)** Two-dimensional continuous parameter scan for CYP2E1 activity (`LI f cyp2e1`) and cirrhosis grade (`f_cirrhosis`). Both parameters were scanned over a continuous range and the pharmacokinetic parameters were calculated. The resulting array was plotted as a heat map. The black lines mark the isoclines. The solid line denotes the isocline for the baseline value, i.e. the pharmacokinetic parameter calculated at `LI_f_cyp2e1` = 1 and `f_cirrhosis` = 0.

To further study the counteracting effects of cirrhosis resulting in reduced liver function and increased CYP2E1 activity, we scanned both parameters (Fig. 8B). Analysis of the isocline representing the simulated MR for the reference values of both parameters (`f_cirrhosis`=0, `LI_f_ycp2e1` =1) reveals that a 2 to 3-fold induction of CYP2E1, typically seen in alcoholics, can compensate for mild liver cirrhosis (`f_cirrhosis`=0.4) in the metabolic phenotyping result. Consequently, special care must be taken when interpreting chlorzoxazone-based metabolic phenotyping results in patients with liver disease.

## 4 DISCUSSION

In this work we developed and validated a physiologically-based pharmacokinetics (PBPK) model for chlorzoxazone used for CYP2E1 metabolic phenotyping. The model was developed and validated on a large database of heterogeneous studies and is freely available in the open standard SBML (Keating et al., 2020).

The model and accurately predicted pharmacokinetic data from various studies. The model successfully described the concentration profiles of chlorzoxazone and its metabolites for various doses. It demonstrated that both changes in CYP2E1 activity and affinity significantly impacted chlorzoxazone pharmacokinetics and metabolic phenotyping results. The model was then used to investigate the effects of alcohol on CYP2E1 and chlorzoxazone pharmacokinetics, finding that changes in CYP2E1 activity due to alcohol consumption could account for the observed alterations in chlorzoxazone pharmacokinetics in alcoholic patients. Furthermore, the model effectively predicted changes in chlorzoxazone pharmacokinetics and metabolic phenotyping due to alcohol induction and withdrawal. Lastly, the model was used to assess the effects of cirrhosis in combination with CYP2E1 induction on liver function. The results indicated that CYP2E1 induction in alcoholic patients with cirrhosis may serve as a compensatory mechanism, partially maintaining metabolic capacity.

The pharmacokinetic data curated for this work displays considerable variability. Most of the studies reported the kinetic data in the form of mean*±*SD or mean*±*SE. However, individual chlorzoxazone time-courses show a large variability in c_max_ and t_max_(de Vries et al., 1994). We fitted the model by minimizing the distance between the time-course data and the model prediction, thus obtaining a parameter set for an average patient. Fitting the model to individual time-courses may allow drawing conclusions about the enzymatic equipment of individual subjects. Microsomal studies report immense interindividual variability. We demonstrated that mapping the microsomal CYP2E1 concentration to the CYP2E1 activity parameter of our model enables predicting the in vivo pharmacokinetics of subject groups (Fig. 6B). It remains to be further investigated whether microsomal enzyme concentration distributions can be used to predict the variability of CYP2E1 kinetics found in clinical pharmacokinetic studies.

We implemented an ethanol pharmacokinetic model capable of describing alcohol metabolism for various doses of oral ethanol administration. Ethanol is primarily metabolized by alcohol dehydrogenase in the liver. Through parameter fitting, we determined the K_M_ value of the enzymatic reaction to be 0.046 µM, which corresponds to the reported k_M_ (0.05 µM) of the ADH1B*1 genotype of alcohol dehydrogenase (Jiang et al., 2020). Although CYP2E1 is known to metabolize ethanol to acetaldehyde with a higher K_M_ than alcohol dehydrogenase, playing a significant role at higher ethanol doses, we did not include this alternative route, as the mono-enzymatic model adequately reproduced the experimental data.

We investigated whether a simple ethanol induction model for CYP2E1 could describe the acceleration of chlorzoxazone metabolism in subjects who recently began drinking, as found by (Oneta et al., 2002). We first determined the half-life of CYP2E1 by comparing our model output to data from several alcohol withdrawal studies. With a first-order decay of CYP2E1 and a half-life of 1.8 d, the data was well-described. However, the induction in subjects who recently started drinking followed more complex dynamics. With the half-life we determined, our model reaches maximum induction after approximately two weeks, while the data did not show saturation even after four weeks of drinking. Three individuals displayed a boost in induction between the third and fourth weeks. This is the only study investigating the dynamics of CYP2E1 induction for individuals who started drinking, and the number of subjects (n=5) is too small to draw definitive conclusions.Additionally, other metabolic state changes in the liver might occur with chronic drinking.

Approximately 90% of drinkers who consume 4 to 5 drinks per day develop steatosis, the initial stage of alcoholic liver disease. Prolonged alcohol consumption can lead to liver inflammation, fibrosis, cirrhosis, and liver cancer (Osna et al., 2017). Depending on its severity, cirrhosis can cause reduced liver function or even liver failure. Our model predicts that in individuals with mild to moderate cirrhotic livers, the baseline level of metabolic function via CYP2E1 can be maintained due to CYP2E1 induction. However, this is even more detrimental in cirrhotic livers, as the intact liver volume is reduced. Although the overall metabolic activity is comparable to that of non-cirrhotic and non-induced livers, the CYP2E1 activity per liver volume is increased. Since cirrhotic livers suffer from severe inflammation, the elevated CYP2E1 activity exposes the remaining tissue to greater oxidative stress, thereby accelerating the progression of the disease.

In conclusion, we developed and validated a PBPK model for CYP2E1 phenotyping using chlorzoxazone.

## AUTHOR CONTRIBUTIONS

JK and MK conceived and designed the study, developed the computational model, curated the data, implemented and performed the analysis, and drafted the manuscript. JG provided valuable support with PK-DB, data curation, and modeling. All authors actively participated in the discussions of the results, contributed to critical revisions of the manuscript, and approved the final version for submission.

## DATA AVAILABILITY STATEMENT

The datasets analyzed for this study can be found in PK-DB available from https://pk-db.com.

## REFERENCES

(2012). LiverTox: Clinical and Research Information on Drug-Induced Liver Injury (Bethesda (MD): National Institute of Diabetes and Digestive and Kidney Diseases)

Bachmann, K. and Sarver, J. G. (1996). Chlorzoxazone as a single sample probe of hepatic CYP2E1 activity in humans. Pharmacology 52, 169–177. doi:10.1159/000139381

Banerjee, A., Kocarek, T. A., and Novak, R. F. (2000). Identification of a ubiquitination-Target/Substrate-interaction domain of cytochrome P-450 (CYP) 2E1. Drug Metabolism and Disposition: The Biological Fate of Chemicals 28, 118–124

Bardag-Gorce, F., Li, J., French, B. A., and French, S. W. (2002). Ethanol withdrawal induced CYP2E1 degradation in vivo, blocked by proteasomal inhibitor PS-341. Free Radical Biology and Medicine 32, 17–21. doi:10.1016/S0891-5849(01)00768-7

Bedada, S. K. and Boga, P. K. (2017). Effect of piperine on CYP2E1 enzyme activity of chlorzoxazone in healthy volunteers. Xenobiotica; the fate of foreign compounds in biological systems 47, 1035–1041. doi:10.1080/00498254.2016.1241450

Bedada, S. K. and Neerati, P. (2016). Resveratrol Pretreatment Affects CYP2E1 Activity of Chlorzoxazone in Healthy Human Volunteers. Phytotherapy research : PTR 30, 463–468. doi:10.1002/ptr.5549

Bedada, S. K. and Neerati, P. (2018). The effect of quercetin on the pharmacokinetics of chlorzoxazone, a CYP2E1 substrate, in healthy subjects. European journal of clinical pharmacology 74, 91–97. doi:10.1007/s00228-017-2345-9

Benowitz, N. L., Peng, M., and Jacob, P., 3rd (2003). Effects of cigarette smoking and carbon monoxide on chlorzoxazone and caffeine metabolism. Clin. Pharmacol. Ther. 74, 468–474. doi:10.1016/j.clpt.2003.07.001

Burckart, G. J., Frye, R. F., Kelly, P., Branch, R. A., Jain, A., Fung, J. J., et al. (1998). Induction of CYP2E1 activity in liver transplant patients as measured by chlorzoxazone 6-hydroxylation. Clinical pharmacology and therapeutics 63, 296–302. doi:10.1016/S0009-9236(98)90161-8

Carriere, V., Goasduff, T., Ratanasavanh, D., Morel, F., Gautier, J. C., Guillouzo, A., et al. (1993). Both cytochromes P450 2E1 and 1A1 are involved in the metabolism of chlorzoxazone. Chemical Research in Toxicology 6, 852–857. doi:10.1021/tx00036a015

Chalasani, N., Gorski, J. C., Asghar, M. S., Asghar, A., Foresman, B., Hall, S. D., et al. (2003). Hepatic cytochrome P450 2E1 activity in nondiabetic patients with nonalcoholic steatohepatitis. Hepatology (Baltimore, Md.) 37, 544–550. doi:10.1053/jhep.2003.50095

Couto, N., Al-Majdoub, Z. M., Achour, B., Wright, P. C., Rostami-Hodjegan, A., and Barber, J. (2019). Quantification of Proteins Involved in Drug Metabolism and Disposition in the Human Liver Using Label-Free Global Proteomics. Molecular Pharmaceutics 16, 632–647. doi:10.1021/acs.molpharmaceut.8b00941

de la Maza, M. P., Hirsch, S., Petermann, M., Suazo, M., Ugarte, G., and Bunout, D. (2000). Changes in microsomal activity in alcoholism and obesity. Alcoholism, clinical and experimental research 24, 605–610. doi:10.1097/00000374-200005000-00004

de Simone, G., Devereux, R. B., Daniels, S. R., Mureddu, G., Roman, M. J., Kimball, T. R., et al. (1997). Stroke volume and cardiac output in normotensive children and adults. Assessment of relations with body size and impact of overweight. Circulation 95, 1837–1843. doi:10.1161/01.cir.95.7.1837

de Vries, J. D., Salphati, L., Horie, S., Becker, C. E., and Hoener, B. A. (1994). Variability in the disposition of chlorzoxazone. Biopharmaceutics &drug disposition 15, 587–597. doi:10.1002/bdd.2510150706

Desiraju, R. K., Renzi, N. L., Nayak, R. K., and Ng, K.-T. (1983). Pharmacokinetics of Chlorzoxazone in Humans. Journal of Pharmaceutical Sciences 72, 991–994. doi:10.1002/jps.2600720905

Dreisbach, A. W., Ferencz, N., Hopkins, N. E., Fuentes, M. G., Rege, A. B., George, W. J., et al. (1995). Urinary excretion of 6-hydroxychlorzoxazone as an index of CYP2E1 activity. Clinical pharmacology and therapeutics 58, 498–505. doi:10.1016/0009-9236(95)90169-8

Eap, C. B., Schnyder, C., Besson, J., Savary, L., and Buclin, T. (1998). Inhibition of CYP2E1 by chlormethiazole as measured by chlorzoxazone pharmacokinetics in patients with alcoholism and in healthy volunteers. Clinical pharmacology and therapeutics 64, 52–57. doi:10.1016/S0009-9236(98)90022-4

Eliasson, E., Johansson, I., and Ingelman-Sundberg, M. (1988). Ligand-dependent maintenance of ethanol-inducible cytochrome P-450 in primary rat hepatocyte cell cultures. Biochemical and Biophysical Research Communications 150, 436–443. doi:10.1016/0006-291X(88)90539-6

Ernstgard, L., Warholm, M., and Johanson, G. (2004). Robustness of chlorzoxazone as an in vivo measure of cytochrome P450 2E1 activity. British Journal of Clinical Pharmacology 58, 190–200. doi:10.1111/j.1365-2125.2004.02132.x

Frye, R. F., Adedoyin, A., Mauro, K., Matzke, G. R., and Branch, R. A. (1998). Use of chlorzoxazone as an in vivo probe of cytochrome P450 2E1: Choice of dose and phenotypic trait measure. Journal of clinical pharmacology 38, 82–89. doi:10.1002/j.1552-4604.1998.tb04381.x

Girre, C., Lucas, D., Hispard, E., Menez, C., Dally, S., and Menez, J. F. (1994). Assessment of cytochrome P4502E1 induction in alcoholic patients by chlorzoxazone pharmacokinetics. Biochemical pharmacology 47, 1503–1508. doi:10.1016/0006-2952(94)90524-x

Grzegorzewski, J., Bartsch, F., Kö ller, A., and Kö nig, M. (2021a). Pharmacokinetics of Caffeine: A Systematic Analysis of Reported Data for Application in Metabolic Phenotyping and Liver Function Testing. Frontiers in Pharmacology 12, 752826. doi:10.3389/fphar.2021.752826

Grzegorzewski, J., Brandhorst, J., Green, K., Eleftheriadou, D., Duport, Y., Barthorscht, F., et al. (2021b). PK-DB: Pharmacokinetics database for individualized and stratified computational modeling. Nucleic Acids Research 49, D1358–D1364. doi:10.1093/nar/gkaa990

He, J., Li, N., Xu, J., Zhu, J., Yu, Y., Chen, X., et al. (2019). An LC-MS/MS Validated Method for Quantification of Chlorzoxazone in Human Plasma and Its Application to a Bioequivalence Study. Journal of chromatographic science 57, 751–757. doi:10.1093/chromsci/bmz052

Herman, I. P. (2016). Physics of the Human Body (Springer)

Hohmann, N., Blank, A., Burhenne, J., Suzuki, Y., Mikus, G., and Haefeli, W. E. (2019). Simultaneous phenotyping of CYP2E1 and CYP3A using oral chlorzoxazone and midazolam microdoses. British journal of clinical pharmacology 85, 2310–2320. doi:10.1111/bcp.14040

Hucka, M., Bergmann, F. T., Chaouiya, C., Dräger, A., Hoops, S., Keating, S. M., et al. (2019). The Systems Biology Markup Language (SBML): Language Specification for Level 3 Version 2 Core Release 2. Journal of Integrative Bioinformatics 16. doi:10.1515/jib-2019-0021

Hukkanen, J., Jacob Iii, P., Peng, M., Dempsey, D., and Benowitz, N. L. (2010). Effects of nicotine on cytochrome P450 2A6 and 2E1 activities. British journal of clinical pharmacology 69, 152–159. doi:10.1111/j.1365-2125.2009.03568.x

ICRP (2002). Basic anatomical and physiological data for use in radiological protection: reference values. a report of age- and gender-related differences in the anatomical and physiological characteristics of reference individuals. icrp publication 89. Annals of the ICRP 32, 5–265

Jiang, Y., Zhang, T., Kusumanchi, P., Han, S., Yang, Z., and Liangpunsakul, S. (2020). Alcohol Metabolizing Enzymes, Microsomal Ethanol Oxidizing System, Cytochrome P450 2E1, Catalase, and Aldehyde Dehydrogenase in Alcohol-Associated Liver Disease. Biomedicines 8, 50. doi:10.3390/biomedicines8030050

Jones, H. and Rowland-Yeo, K. (2013). Basic concepts in physiologically based pharmacokinetic modeling in drug discovery and development. CPT: pharmacometrics &systems pharmacology 2, e63. doi:10.1038/psp.2013.41

Keating, S. M., Waltemath, D., König, M., Zhang, F., Dräger, A., Chaouiya, C., et al. (2020). SBML Level 3: An extensible format for the exchange and reuse of biological models. Molecular Systems Biology 16. doi:10.15252/msb.20199110

Kharasch, E. D., Thummel, K. E., Mhyre, J., and Lillibridge, J. H. (1993). Single-dose disulfiram inhibition of chlorzoxazone metabolism: A clinical probe for P450 2E1. Clinical pharmacology and therapeutics 53, 643–650. doi:10.1038/clpt.1993.85

Kö ller, A., Grzegorzewski, J., and Kö nig, M. (2021a). Physiologically based modeling of the effect of physiological and anthropometric variability on indocyanine green based liver function tests. Frontiers in physiology, 2043

Kö ller, A., Grzegorzewski, J., Tautenhahn, H.-M., and Kö nig, M. (2021b). Prediction of survival after partial hepatectomy using a physiologically based pharmacokinetic model of indocyanine green liver function tests. Frontiers in physiology, 1975

[Dataset] Kö nig, M. (2021a). Sbmlsim: SBML simulation made easy. Zenodo. doi:10.5281/ZENODO.5531088

[Dataset] Kö nig, M. (2021b). Sbmlutils: Python utilities for SBML. Zenodo. doi:10.5281/ZENODO.5546603

Kö nig, M., Dräger, A., and Holzhü tter, H.-G. (2012). CySBML: A Cytoscape plugin for SBML. Bioinformatics 28, 2402–2403. doi:10.1093/bioinformatics/bts432

[Dataset] Kö nig, M. and Rodriguez, N. (2019). Matthiaskoenig/cy3sbml: Cy3sbml-v0.3.0 - SBML for Cytoscape. Zenodo. doi:10.5281/ZENODO.3451319

[Dataset] Kü ttner, J. and König, M. (2023). Physiologically based pharmacokinetic (PBPK) model of chlorzoxazone. Zenodo. doi:10.5281/ZENODO.7821956

Liangpunsakul, S., Kolwankar, D., Pinto, A., Gorski, J. C., Hall, S. D., and Chalasani, N. (2005). Activity of CYP2E1 and CYP3A enzymes in adults with moderate alcohol consumption: A comparison with nonalcoholics. Hepatology (Baltimore, Md.) 41, 1144–1150. doi:10.1002/hep.20673

Lucas, D., Berthou, F., Girre, C., Poitrenaud, F., and Ménez, J.-F. (1993). High-performance liquid chromatographic determination of chlorzoxazone and 6-hydroxychlorzoxazone in serum: A tool for indirect evaluation of cytochrome P4502E1 activity in humans. Journal of Chromatography B: Biomedical Sciences and Applications 622, 79–86. doi:10.1016/0378-4347(93)80252-Y

Lucas, D., Ferrara, R., Gonzalez, E., Bodenez, P., Albores, A., Manno, M., et al. (1999). Chlorzoxazone, a selective probe for phenotyping CYP2E1 in humans. Pharmacogenetics 9, 377–388. doi:10.1097/00008571-199906000-00013

Lucas, D., Ménez, C., Girre, C., Bodénez, P., Hispard, E., and Ménez, J. F. (1995). Decrease in cytochrome P4502E1 as assessed by the rate of chlorzoxazone hydroxylation in alcoholics during the withdrawal phase. Alcoholism, clinical and experimental research 19, 362–366. doi:10.1111/j.1530-0277.1995.tb01516.x

Mishin, V. M., Rosman, A. S., Basu, P., Kessova, I., Oneta, C. M., and Lieber, C. S. (1998). Chlorzoxazone pharmacokinetics as a marker of hepatic cytochrome P4502E1 in humans. The American Journal of Gastroenterology 93, 2154–2161. doi:10.1111/j.1572-0241.1998.00612.x

Oneta, C. M., Lieber, C. S., Li, J., Rü ttimann, S., Schmid, B., Lattmann, J., et al. (2002). Dynamics of cytochrome P4502E1 activity in man: Induction by ethanol and disappearance during withdrawal phase. Journal of hepatology 36, 47–52. doi:10.1016/s0168-8278(01)00223-9

Ono, S., Hatanaka, T., Hotta, H., Tsutsui, M., Satoh, T., and Gonzalez, F. J. (1995). Chlorzoxazone is metabolized by human CYP1A2 as well as by human CYP2E1. Pharmacogenetics 5, 143–150. doi:10.1097/00008571-199506000-00002

Orellana, M., Rodrigo, R., Varela, N., Araya, J., Poniachik, J., Csendes, A., et al. (2006). Relationship between in vivo chlorzoxazone hydroxylation, hepatic cytochrome P450 2E1 content and liver injury in obese non-alcoholic fatty liver disease patients. Hepatology Research 34, 57–63. doi:10.1016/j.hepres.2005.10.001

O’Shea, D., Davis, S. N., Kim, R. B., and Wilkinson, G. R. (1994). Effect of fasting and obesity in humans on the 6-hydroxylation of chlorzoxazone: A putative probe of CYP2E1 activity. Clinical pharmacology and therapeutics 56, 359–367. doi:10.1038/clpt.1994.150

Osna, N. A., Donohue, T. M., and Kharbanda, K. K. (2017). Alcoholic Liver Disease: Pathogenesis and Current Management. Alcohol Research: Current Reviews 38, 147–161

Park, J.-Y., Kim, K.-A., Park, P.-W., and Ha, J.-M. (2006). Effect of high-dose aspirin on CYP2E1 activity in healthy subjects measured using chlorzoxazone as a probe. Journal of clinical pharmacology 46, 109–114. doi:10.1177/0091270005282635

Rajnarayana, K., Venkatesham, A., Nagulu, M., Srinivas, M., and Krishna, D. R. (2008). Influence of diosmin pretreatment on the pharmacokinetics of chlorzoxazone in healthy male volunteers. Drug metabolism and drug interactions 23, 311–321. doi:10.1515/dmdi.2008.23.3-4.311

Raucy, J. L., Kraner, J. C., and Lasker, J. M. (1993). Bioactivation of Halogenated Hydrocarbons by Cytochrome P4502E1. Critical Reviews in Toxicology 23, 1–20. doi:10.3109/10408449309104072

Roberts, B. J., Song, B.-J., Soh, Y., Park, S. S., and Shoaf, S. E. (1995). Ethanol Induces CYP2E1 by Protein Stabilization. Journal of Biological Chemistry 270, 29632–29635. doi:10.1074/jbc.270.50.29632

Somogyi, E. T., Bouteiller, J.-M., Glazier, J. A., Kö nig, M., Medley, J. K., Swat, M. H., et al. (2015). libRoadRunner: A high performance SBML simulation and analysis library. Bioinformatics 31, 3315– 3321. doi:10.1093/bioinformatics/btv363

Song, B. J., Veech, R. L., Park, S. S., Gelboin, H. V., and Gonzalez, F. J. (1989). Induction of Rat Hepatic N-Nitrosodimethylamine Demethylase by Acetone Is Due to Protein Stabilization. Journal of Biological Chemistry 264, 3568–3572. doi:10.1016/S0021-9258(18)94103-7

Takahashi, T., Lasker, J. M., Rosman, A. S., and Lieber, C. S. (1993). Induction of cytochrome P-4502E1 in the human liver by ethanol is caused by a corresponding increase in encoding messenger RNA. Hepatology 17, 236–245. doi:10.1002/hep.1840170213

Tanaka, E., Terada, M., and Misawa, S. (2000). Cytochrome P450 2E1: Its clinical and toxicological role. Journal of Clinical Pharmacy and Therapeutics 25, 165–175. doi:10.1046/j.1365-2710.2000.00282.x

Urso, R., Blardi, P., and Giorgi, G. (2002). A short introduction to pharmacokinetics. European review for medical and pharmacological sciences 6, 33–44

Vander, J. S., A. (2001). Human physiology: The mechanisms of body function. McGraw-Hill higher education

Vesell, E. S., Seaton, T. D., and A-Rahim, Y. I. (1995). Studies on interindividual variations of CYP2E1 using chlorzoxazone as an in vivo probe. Pharmacogenetics 5, 53–57. doi:10.1097/00008571-199502000-00007

Virtanen, P., Gommers, R., Oliphant, T. E., Haberland, M., Reddy, T., Cournapeau, D., et al. (2020). SciPy 1.0: Fundamental Algorithms for Scientific Computing in Python. Nature Methods 17, 261–272. doi:10.1038/s41592-019-0686-2

Wang, Z., Hall, S. D., Maya, J. F., Li, L., Asghar, A., and Gorski, J. C. (2003). Diabetes mellitus increases the in vivo activity of cytochrome P450 2E1 in humans. British journal of clinical pharmacology 55, 77–85. doi:10.1046/j.1365-2125.2003.01731.x

Welsh, C., Xu, J., Smith, L., König, M., Choi, K., and Sauro, H. M. (2023). libRoadRunner 2.0: A high performance SBML simulation and analysis library. Bioinformatics 39, btac770. doi:10.1093/bioinformatics/btac770

Wilkinson, P. K., Sedman, A. J., Sakmar, E., Kay, D. R., and Wagner, J. G. (1977). Pharmacokinetics of ethanol after oral administration in the fasting state. Journal of Pharmacokinetics and Biopharmaceutics 5, 207–224. doi:10.1007/B01065396

Witt, L., Suzuki, Y., Hohmann, N., Mikus, G., Haefeli, W. E., and Burhenne, J. (2016). Ultrasensitive quantification of the CYP2E1 probe chlorzoxazone and its main metabolite 6-hydroxychlorzoxazone in human plasma using ultra performance liquid chromatography coupled to tandem mass spectrometry after chlorzoxazone microdosing. Journal of chromatography. B, Analytical technologies in the biomedical and life sciences 1027, 207–213. doi:10.1016/j.jchromb.2016.05.049

Yamamura, Y., Koyama, N., and Umehara, K. (2015). Comprehensive kinetic analysis and influence of reaction components for chlorzoxazone 6-hydroxylation in human liver microsomes with CYP antibodies. Xenobiotica; the fate of foreign compounds in biological systems 45, 353–360. doi:10.3109/00498254.2014.985760

Zhang, H.-F., Wang, H.-H., Gao, N., Wei, J.-Y., Tian, X., Zhao, Y., et al. (2016). Physiological Content and Intrinsic Activities of 10 Cytochrome P450 Isoforms in Human Normal Liver Microsomes. The Journal of Pharmacology and Experimental Therapeutics 358, 83–93. doi:10.1124/jpet.116.233635

